# Genomic animal models quantify the heritable components of spatiotemporally structured phenotypic variation in a dispersive cryptobenthic marine fish

**DOI:** 10.1101/2024.11.25.625127

**Authors:** Joshua A. Thia, J. David Aguirre, Jennifer Evans, James Hereward, Libby Liggins, Will Figueira, Cynthia Riginos, Katrina McGuigan

## Abstract

In highly dispersive taxa, gene flow can homogenise genetic variation over broad spatial and temporal scales. However, phenotypic differentiation can still arise when heritable genetic variation is available to selection and (or) when organisms exhibit plasticity. Disentangling heritable versus non-heritable components of variation is possible using genomic animal models, but this approach has been underused in wild organisms.

We investigated head shape variation in the marine cryptobenthic fish *Bathygobius cocosensis*, a highly dispersive species occupying heterogeneous environments.

Head shape, a diet-related trait, is spatially structured in Australian populations. We curated a unique dataset comprising genotypic, phenotypic, and environmental variation to: (1) contrast the scales over which genetic and phenotypic variation are structured; and (2) quantify heritable versus non-heritable contributions to head shape variation using genomic animal models and comparisons between *F*_ST_, *P*_ST_, and *Q*_ST_.

At macrogeographic scales (hundreds of kilometres) and over time (2 years), phenotypic differentiation (0.0001 ≤ *P*_ST_ ≤ 0.47) exceeded both genomic and additive genetic differentiation (0.001 ≤ *F*_ST_≤ 0.004 and 0.00001 ≤ *Q*_ST_ ≤ 0.004, respectively). This suggests head shape variation may primarily reflect (non-heritable) plastic responses, rather than (heritable) adaptive genetic changes in mean head shape. At microgeographic scales, associations between head shape and tide pool variables were temporally variable, indicating that local environmental conditions may structure phenotypes within populations.

Overall, this study highlights the value of genomic animal models for disentangling heritable and non-heritable components of phenotypic variation in dispersive marine taxa. It provides an integrative analysis linking genetic, phenotypic, and environmental variation in a cryptobenthic fish, highlighting how phenotypic divergence can emerge without strong genetic differentiation, and challenging the common assumption that phenotypic divergence necessarily implies adaptation.

## INTRODUCTION

In species with large population sizes and high dispersal potential, extensive gene flow is expected to homogenise allele frequencies (Waples, 1998), resulting in weak neutral genetic differentiation across space and time. Paradoxically, such systems often exhibit substantial spatial variation in ecologically important phenotypic traits, even across environmental gradients that are well within dispersal distances (Tigano & Friesen, 2016). Theoretical developments alongside these key empirical examples have shifted the perception of the potential for local adaptation to contribute trait divergence even under high gene flow (Yeaman & Whitlock, 2011; Yeaman, 2022) However, phenotypic differentiation between environments can arise through divergent evolution of trait mean (adaptation), and (or) through environmentally induced shifts in trait expression without genetic change (phenotypic plasticity) (Hoffmann *et al*., 2005; Lind *et al*., 2011; Moody *et al*., 2015).

Disentangling causes of phenotypic heterogeneity in the wild remains extremely challenging for multiple reasons. Difficulties in accurately measuring natural selection (Kingsolver et al. 2012; Shaw 2019), and the necessity of manipulative experiments to identify specific causal environmental factors, limit our understanding of adaptive

processes (Wadgymar *et al*., 2026). Because local adaptation is defined by divergence in allele frequency in the direction of selection, the accessibility of genomic tools suggests that we should be able to readily test whether observed phenotypic differences are due to adaptive evolution through association tests. However, for polygenic traits, signatures of adaptive evolution may remain difficult to detect (but for examples, see: Babin et al., 2017; Sella & Barton, 2019; Rey et al., 2020; Tepolt et al., 2021). Phenotypic plasticity has been considered the most parsimonious explanation when adaptive (genotypic) divergence is not supported (Merilä & Hendry, 2014). However, the low power of many studies of wild organisms means that the absence of a signal in genomic association studies (Santure & Garant, 2018; Duntsch *et al*., 2020; Lasky *et al*., 2022), and therefore, the importance of adaptive plasticity, should be interpreted cautiously.

A long-standing, but underutilised approach for resolving the importance of heritable adaptive divergence among populations is to compare the differentiation at neutral genetic markers (*F*_ST_) to that of additive genetic variance in ecologically important, putatively adaptive traits (*Q*_ST_) (Merilä & Crnokrak, 2001; Chenoweth & Blows, 2008). Under neutral evolution, the expectation is that *F*_ST_ = *Q*_ST_, whereas under adaptive evolution, *F*_ST_ < *Q*_ST_ (Merilä & Crnokrak, 2001). One reason why this approach is not more commonly used is that partitioning of phenotypic variation to the heritable (additive genetic) component has historically depended on controlled breeding or multigenerational data from natural populations to determine familial relationships among individuals, both implausible approaches for many taxa (Merilä & Crnokrak, 2001). When additive genetic components cannot be measured directly, studies have often used the analogue *P*_ST_, which quantifies phenotypic differentiation among populations based on the partitioning of total phenotypic variance. However, the interpretation of *P*_ST_ as a proxy for additive genetic differentiation depends on assumptions about heritability (Leinonen *et al*., 2006; Raeymaekers *et al*., 2007; Brommer, 2011).

Genomic animal models provide a means to obtain estimates of additive genetic variance in wild organisms where controlled breeding experiments or intensive multigenerational study of natural populations are not feasible (Kruuk, 2004; Garant & Kruuk, 2005; Schmidt *et al*., 2023). These models extend classical quantitative genetic approaches by estimating genetic similarity among individuals directly from genome-wide markers rather than from pedigrees (Kruuk, 2004; Frentiu *et al*., 2008; Wilson *et al*., 2010; Bérénos *et al*., 2014). Under this genomic approach, the degree of relatedness among all pairs of individuals is characterised directly by their proportion of shared alleles (Porth *et al*., 2015; Gervais *et al*., 2019; James *et al*., 2022). Despite their potential value, genomic animal models have been largely underused in the study of wild organisms (Schmidt *et al*., 2023), but have been applied to, for instance, natural populations of hihi bird (Duntsch *et al*., 2020), soay sheep (James *et al*., 2022), roe deer (Gervais *et al*., 2019), and stickleback fish (Fraimout *et al*., 2024). Greater exploitation of genomic animal models could widen the scope for evolutionary studies, both in terms of estimating additive genetic variance in natural populations, but also in testing hypotheses of adaptive genetic divergence.

Intertidal fishes provide a compelling system for studying phenotypic differentiation across different spatiotemporal scales. Many are so-called ‘cryptobenthic fishes’, a diverse group of small, often camouflaged, blennies and gobies that inhabit the benthos of marine habitats (Goatley & Brandl, 2017; Brandl et al., 2018). Their inconspicuous morphology and (or) behaviour make them difficult to study, and as a result, their evolutionary ecology and contribution to marine ecosystem dynamics remain poorly understood (Goatley & Brandl, 2017; Brandl et al., 2018). Many cryptobenthic fishes inhabiting reefs and rocky intertidal zones exhibit marked shifts in dispersal scale across their life cycle (Mukai et al., 2009; Čekovská et al., 2020; Sefc et al., 2020). Species with a prolonged pelagic larval phase may disperse over long distances, potentially associated with markedly diverse environmental conditions between populations. After settlement, these fish can encounter environmental heterogeneity on the microgeographic scale across the reef or intertidal habitat patches they occupy. In rocky intertidal habitats, local environmental conditions can vary considerably across the tidal gradient. Such within-population environmental heterogeneity can drive phenotype divergences on microgeographic spatial scales, as seen in mussels, barnacles and gastropods (Schmidt & Rand, 1999; Westram et al., 2014; Richards et al., 2025).

Our focal species in this present study is *Bathygobius cocosensis* (Bleeker, 1854: Perciformes; Gobiidae). Characteristic of cryptobenthic fish, *B. cocosensis* is small (adults are typically <60 mm) and inhabits shallow marine and intertidal benthic environments across the Indo Pacific (Griffiths, 2003b; Mukai et al., 2009). Prior work on Australian populations of *B. cocosensis* has analysed patterns of phenotypic variation in head shape (Malard *et al*., 2016; Thia *et al*., 2021), given the ecological importance of head shape in fishes for prey acquisition (for example: Marcil et al., 2006; Vera-Duarte et al., 2017). Head shape phenotypes in *B. cocosensis* are structured across sites over hundreds of kilometres of coastline, suggesting divergent selection pressures at macrogeographic scales (Thia et al., 2021).

Differences in head shape have also been reported at scales of tens of meters along a tidal elevation gradient within a single site, indicating that microhabitat variation may drive adaptive responses within populations on microgeographic scales (Malard et al., 2016). The phenotypic differentiation of head shape in *B. cocosensis* at both large and small spatial scales is particularly intriguing given the very weak population genetic structure among populations in eastern Australia (Thia et al., 2021). Reconciling high dispersal potential and low genetic structure with moderate phenotypic divergence therefore remains a key question in this species.

In the present work, we investigate the spatiotemporal variability of genetic and phenotypic structure in *B. cocosensis* and demonstrate the utility of genomic animal models to estimate quantitative genetic variation in wild organisms. We curated a unique dataset in which *B. cocosensis* individuals sampled over multiple years (distinct cohorts), and from environmentally distinct sites, were phenotyped using geometric morphometrics and genotyped using reduced-representation sequencing (RAD-seq). Across sites (macrogeographic scale) and over multiple years, we quantified structure in: (1) genotypes (*F*_ST_); (2) head shape phenotypes (*P*_ST_); and (3) heritable components of head shape (*Q*_ST_). Within a single site (microgeographic scale) and over multiple years, we tested for local structuring associated with different microhabitat variables. Overall, we show structuring of phenotypic variation exceeds that of genetic variation. Whilst some components of head shape appeared heritable, as estimated from genomic animal models, there was no evidence that additive genetic variance was structured among sites or years. Our results might therefore suggest a greater importance of plastic responses in shaping the distribution of head shape in *B. cocosensis*.

## MATERIALS AND METHODS

### Sample collection and study sites

Our sampling design aimed to capture distinct environments on a macrogeographic scale as well as potential temporal variation between different generations. Our two study sites were located on Australia’s eastern coastline: Hastings Point (S28.36° E153.58°, New South Wales) and Heron Island (S23.44° E151.91°, Queensland), which respectively have temperate-subtropical and tropical climates. Hastings Point (Figure 1a) is a rocky headland with a topologically complex intertidal zone with discrete pools that the gobies retreat to during low tide. On Heron Island (Figure 1b), situated ∼80 km from the mainland at the southern limit of Great Barrier Reef coral islands, the intertidal gradient is shallow, tide pools absent, and *B. cocosensis* retreat to the subtidal areas during low tide.

**Figure 1.**
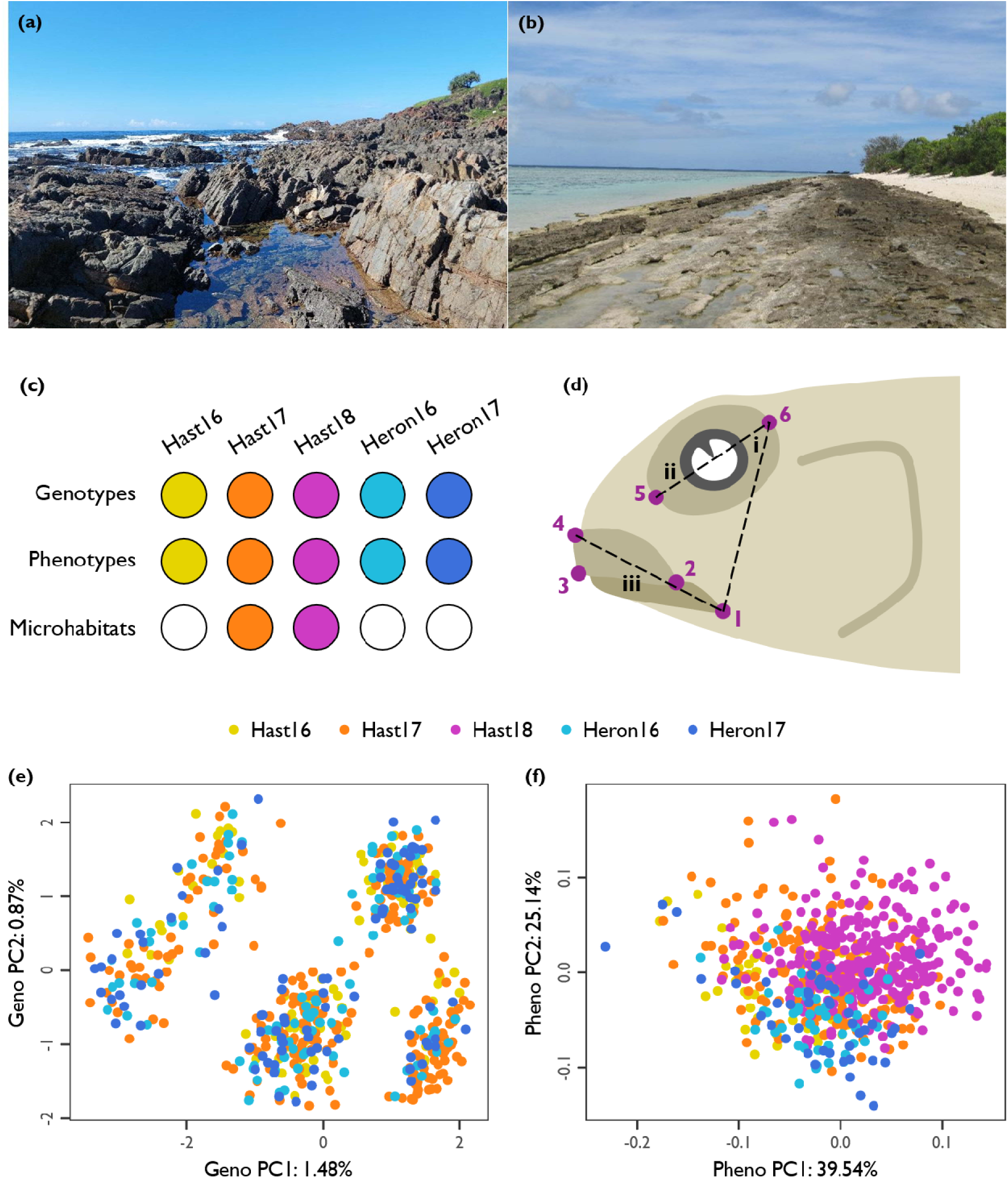
Overview of our study of macro- and microgeographic structure and heritability in *B. cocosensis*. (a) Our first study site, Hastings Point. Photo credit: Joshua Thia. (b) Our second study site, Heron Island. Photo credit: Joshua Thia. (c) Data collected for our study. Filled circles indicate the types of data (rows) collected for each spatiotemporally sampled population (columns). (d) Schematic of 2D landmarks (numbered points) and classic morphometric measurements (Roman numerals) describing head shape morphology. Classical measurements describe eye angle (i), eye size (ii) and mouth size (iii). (e) PCA of genotypic variation. (f) PCA of phenotypic variation. (e,f) The *x*-axis is PC1, and the *y*-axis is PC2, with percentages in axis labels indicating the amount of variation explained by each PC. Points represent individual fish, coloured by population (see legend).

Annual collections of *B. cocosensis* were made between 2016 and 2018. At Hastings Point, sampling occurred on the 25^th^ of February 2016, 24^th^ and 25^th^ of February 2017, and 1^st^ and 2^nd^ of March 2018. At Heron Island, sampling occurred on 14^th^ of January 2016 and 11^th^ of January 2017. Fish were caught using hand nets and euthanised with 500 mg/mL eugenol in seawater following ethically approved protocols (University of Sydney #2015/384; University of Queensland #SBS/221/15/HSF). Fish of all ages were included in this study, ranging from young juveniles (∼6 mm) to large adults (∼60 mm). In this study, we refer to a ‘population’ as fish collected from the same site (Hastings Point or Heron Island) in the same year (2016, 2017 and 2018). We use this spatiotemporal definition of populations because of the distinct environmental conditions at the two sites (detailed above), and because the lifespan of *B. cocosensis* is estimated to be less than one year (Thia *et al*., 2021), suggesting that each annual collection largely comprises a different cohort. Where possible, we collected genotype, phenotype, and microhabitat data for each fish (see Figure 1c for sampling design).

### Genotype data

Genome-wide SNP genotypes were obtained to quantify population genetic structure and estimate the additive genetic relationships between individuals. We used a custom RAD-seq (ddRAD) procedure with the *Eco*RI and *Sph*I restriction enzymes (see Supplementary Methods, ‘Custom ddRAD protocol’). The prepared RAD-seq libraries were sequenced on an Illumina HighSeq X platform by AnnoRoad Gene Technology (Beijing, China). Reads were demultiplexed using the *process_radtags* function from *stacks2* v2.55 (Catchen *et al*., 2013; Rochette *et al*., 2019). Reads were quality trimmed using *fastp* v0.12.4 (Chen *et al*., 2018) and were truncated to a final length of 100 bp. There was a mean of 5,201,124 trimmed reads across samples, with a range of 10,928–21,942,052 reads.

RAD tags were assembled *de novo* using the *stacks2* functions *ustacks*, *cstacks*, *sstacks*, *tsv2bam*, and *gstacks*; the *populations* function was used to call SNPs. *Vcftools* v0.1.16 (Danecek *et al*., 2011) was used to perform an initial filtering to remove SNPs with minor allele counts (MAC) <3, genotype qualities <15, and missing data rates >50%. RAD tags were mapped against the *B. cocosensis* mitogenome (Evans *et al*., 2018) to identify sequences that were not derived from the nuclear genome so they could be filtered out downstream: we used *seqtk* v1.3-r106 (Li, 2019) to reformat RAD tag sequences, *bwa mem* v0.7.17 (Li & Durbin, 2009) for mapping, and *samtools* v1.7 (Li *et al*., 2009) for filtering alignments.

SNPs were imported into R for further curation. We used a stepwise procedure that accounted for the multifaceted nature of missing data: samples and loci can vary in their proportions of missing genotypes, and even loci within RAD tags may vary in missing data rates. SNPs derived from the mitogenome were removed. Any samples with >50% missing data were removed because these disproportionately affected rates of missing data across SNPs. We imputed genotypes using the R package, *polyRAD* v2.0.0 (Clark *et al*., 2019), which allowed us retain SNP loci with low coverage. Next, we removed SNPs with missing data >40%, SNPs that were not observed in at least 5 individuals, and SNPs with evidence of heterozygosity excess.

Our final working genotypic dataset comprised 542 samples: 97 and 242 fish from Hastings Point in 2016 and 2017 (respectively), and 100 and 103 fish from Heron Island in 2016 and 2017 (respectively). We used two different SNP sets. A full SNP set comprised 4,414 RAD tags and 31,095 SNPs, was used to estimate the genomic relatedness matrix, which was used to estimate trait SNP heritability. A thinned SNP set, obtained by randomly sampling one SNP per RAD tag, was used for quantifying genetic differentiation among populations.

#### Phenotype data

We measured head shape, a putatively important ecological trait in *B. cocosensis* (Malard *et al*., 2016; Thia *et al*., 2021), across 658 fish. Landmarks capturing variability in eye and mouth shape, as described elsewhere (Thia *et al*., 2021), were placed on 2D images of fish heads using *tpsdig2* (Rohlf, 2015). Landmarks were imported into R and subjected to general Procrustes alignment using the *geomorph* package (Adams & Otárola-Castillo, 2013; Adams *et al*., 2016). The centroid size, which describes the spread of landmarks for an individual (Bookstein, 1997), was retained as a measure of head size.

We characterised head shape using a set of principal components (PCs), reducing dimensionality to simplify downstream analyses. We obtained PCs by applying R’s *prcomp* function to the covariance matrix of aligned head shape landmarks. The leading phenotypic PC axes explained 39.54%, 25.14%, and 15.11% of the total variation in head shape, respectively (Supplementary Figure S1), and 79.79% in total. All downstream analyses of phenotype used these leading 3 axes. To aid in the interpretation of our focal phenotypic PCs, we also obtained from the aligned landmarks classical morphometric measurements: eye size, eye angle, and mouth size (Figure 1b; Supplementary Table S1).

#### Microhabitat data

We collected microhabitat data at Hastings Point to perform fine scale phenotype–environment associations within local populations. We recorded four microhabitat variables that described continuous variation in tide pool environments: inundation, rugosity, volume, and depth. Inundation was the time for tide pools to be inundated on the incoming tide (minutes). Pools in the low intertidal zone had shorter inundation times relative to pools in the high intertidal zone. Rugosity measured the structural complexity of tide pools. The longest axis for each focal tide pool was identified, and two lengths were obtained using a tape measure (meters): the direct length, which was the distance end to end, and the conforming length, which tracked the topography of the substrate surface. The rugosity score was calculated as the ratio of the direct to conforming length (Griffiths *et al*., 2003; Willis *et al*., 2005).

Volume of focal tide pools was measured as the amount of water (litres) they contained at low tide. A bilge pump was used to empty tide pools; the water was stored and measured in 20 L buckets and returned to tide pools after measuring. Depth (cm) was measured as the distance from the deepest point of focal tide pools to the water surface at low tide. In total, we had measurements for 10 and 11 tide pools with at least four representative fish from collections at Hastings Point in 2017 (160 fish) and 2018 (272 fish), respectively. All tide pools sampled in 2017 were also sampled in 2018.

#### Macrogeographic structuring of genotypic and phenotypic variation

We investigated the differentiation in genotypes and phenotypes measured across the two sites sampled in each year. First, we quantified pairwise genetic structure between populations as *F*_ST_ following (Weir & Cockerham, 1984) using *genomalicious* (Thia, 2024). These *F*_ST_ calculations used all 542 fish in our final thinned genotype dataset, which comprised a total of 542 fish from Hastings Point and Heron Island in 2016 and 2017. To test the null hypothesis of no structure (*F*_ST_= 0), a null distribution was generated using 250 permutations of the data, where each sample was randomly allocated to a population and *F*_ST_ was recalculated. We complimented interpretation of *F*_ST_with a principal components analysis (PCA) to assess patterns of genome-wide population structure (Supplementary Figure S2). PCA was performed with R’s *prcomp* function on the genotype covariance matrix.

Next, we analysed phenotypic structure for all populations. The data for phenotypic structure included a total of 658 fish obtained from Hastings Point 2016, 2017 and 2018, and from Heron Island in 2016 and 2017. For the 3 leading phenotypic PCs, we quantified pairwise *P*_ST_ among population pairs. Historically, *P*_ST_ has been used as a proxy for *Q*_ST_ (Spitze, 1993) when experimental designs do not afford estimation of additive genetic effects. However, inference of adaptive genetic structuring from *P*_ST_ needs to be made cautiously in such circumstances where additive genetic effects are not directly quantifiable (Merilä & Crnokrak, 2001; Leinonen *et al*., 2006; Raeymaekers *et al*., 2007). Here, we instead use *P*_ST_ as a more general estimate of the total phenotypic differentiation between populations that could arise from heritable and non-heritable phenotypic differences. Our formulation of *P*_ST_ follows Leinonen et al. (2011):

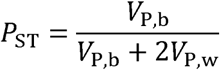

Here, *V*_P,b_and *V*_P,w_ are the phenotypic variances between and within populations, respectively. We obtained these variance components for each phenotypic PC axis and population pair by fitting linear models with *lm*, taking the form:

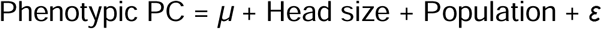

Here, the response, phenotypic PC1, PC2 or PC3, was fit as a function of the fixed effects of ‘Head size’ (continuous log10-transformed centroid size standardised to mean of 0 and standard deviation of 1) and ‘Population’ (categorical population ID). Interpretation of the interaction term was precluded by slight differences in the head size range across populations, and the interaction was therefore not fit. Parameters *µ* and *ε* are the mean and residual error terms, respectively. From these model fits, we retained the least squares mean estimates of the ‘Population’ to estimate *V*_P,b_, and *e* to estimate *V*_P,w_. To test the null hypothesis of no structure (*P*_ST_ = 0), a null distribution was generated using 250 permutations of the data, where each sample was randomly allocated to a population, models were re-fitted, and *P*_ST_ was recalculated.

To gain further insight into the contribution of site (Hastings Point and Heron Island) and sampling year to variation in head shape, we used a multivariate analysis of variance (MANOVA) to partition variation to these factors. Using R’s *manova* function, we fit the model:

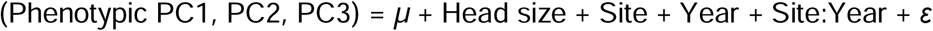

Here, the response, the matrix of our focal phenotypic axes, was fitted as a function of the continuous fixed effects of ‘Head size’ (log10 standardised, as described above), and the categorical fixed effects of sampling ‘Site’ and ‘Year’. Terms *µ* and *ε* are the global mean and residuals, respectively. We determined the variation in our focal phenotypic axes attributed to each of the predictor variables by dividing the trace of each respective sums-of-squares and cross-products matrix by the total variance (sum of traces across all predictor variables).

### Heritability and macrogeographic structuring of additive genetic variance in head shape

We used quantitative genetic approaches to assess the heritability of head shape phenotypes and the potential role of adaptive divergence in structuring heritable variation. First, we estimated the additive genetic variance (*V*_A_) of head shape using genomic animal models (Kruuk, 2004; Garant & Kruuk, 2005; Frentiu *et al*., 2008). The genomic relatedness matrix, **A**_G_, described the realised additive genetic relationships between individuals in the full genotype dataset. We used all 542 genotyped fish from Hastings Point and Heron Island in 2016 and 2017 to improve estimation, but subset the matrix later to include only those with measured head shape phenotypes (see below).

We used the *A.mat* function from R’s *sommer* package (Covarrubias-Pazaran, 2016) to obtain **A**_G_. Kin structure in **A**_G_ can influence estimates of additive genetic variance from genomic animal models. The presence of too many close relatives can bias estimates of *V*_A_ from genomic relatedness matrices (Yang *et al*., 2017; Huang *et al*., 2025). But in contrast, a lack of variation in relatedness can reduce the power to detect and accurately estimate *V*_A_, which is a concern when sampling wild organisms that have large population sizes (Frentiu *et al*., 2008; Fraimout *et al*., 2024). The average relatedness of between pairs of *B. cocosensis* was near zero, with a few pairs of individuals approaching the level of relatedness expected for first cousins (Supplementary Figure S3). Hence, our dataset likely comprises mostly (if not entirely) unrelated individuals.

Next, we subset the relatedness matrix to the 298 fish for which we had both head shape phenotypes and genotypes. We then obtained the nearest positive-definite approximation of this subset matrix using the *nearPD* function from the *Matrix* package (Bates *et al*., 2025). We then fit genomic animal models using *sommer*’s *mmes* function:

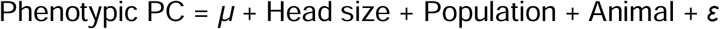

Here, the response, the phenotypic PC, was fit as a function of the fixed effects ‘Head size’ (continuous log10-transformed centroid size standardised to mean of 0 and standard deviation of 1) and ‘Population’ (categorical population ID), with global mean and error represented by *µ* and *ε*, respectively. The term ‘Animal’ represents the random intercept for each individual, corresponding to their additive genetic breeding value, with covariance structure defined by **A**_G_. We assessed the statistical support for non-zero additive genetic variance by fitting a null model that omitted the additive genetic variance (‘Animal’) component and compared this to the full model using a likelihood ratio test (LRT: implemented using *sommer*’s *anova.mmes* function).

To place estimates of *V*_A_ on a standardised scale for interpretation, we calculated narrow-sense heritability as ℎ^2^ = *V*_A_/*V*_P_. To ensure unbiased ℎ^2^ estimates (Wilson *et al*., 2010), *V*_P_ was calculated from both residual variance (*V*_E_) and the variance accounted for by fixed effects (*V*_F_), as detailed in de Villemereuil et al. (2018). Confidence intervals of variance components and heritability were calculated using restricted maximum likelihood multivariate normal (REML-MVN) approach (Houle & Meyer, 2015; Dugand *et al*., 2025). Briefly, this approach involved obtaining 100,000 random samples of *V*_A_ and *V*_E_ using draws from a normal distribution, *N* ∼ (*θ*^, *V*), where *θ*^ was the observed (REML) parameter estimate, and *V* was the asymptotic variance estimate (Fisher Information matrix) obtained from the model. For each random sample, we calculated ℎ^2^ based on the drawn *V*_A_ and *V*_E_ and the observed *V*_F_. Note, this means that *V*_F_ was constant across all random samples.

We then quantified the pairwise additive genetic differentiation between populations as *Q*_ST_, following the Prout-Barker-Spitze formulation (Prout & Barker, 1993; Spitze, 1993), which is the appropriate contrast to Weir and Cockerham’s *F*_ST_ (Liu & Edge, 2025). We calculated *Q*_ST_ using code adapted from Liu & Edge (2025), whereby:

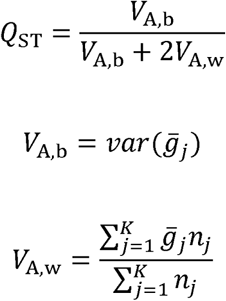

Here, *Q*_ST_ is a function of the additive genetic variances between and within populations, *V*_G,b_ and *V*_G,w_, respectively. *V*_G,b_ is calculated as variance of mean breeding values for each focal phenotypic PC within each *j*^th^ population, *g*_*j*_, and *V*_G,w_ as the weighted average of the variance in breeding values within each *j*^th^ population of sample size *n*_*j*_ across all *K* populations. Breeding values were obtained from the genomic animal models.

Statistical support for adaptive genetic divergence among populations in heritable head shape variation was evaluated using a chi-square test. Under a Lewontin-Krakauer distribution, *Q*_ST_ is expected to be distributed as:

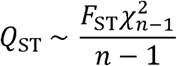

Here, *F*_ST_is the genome-wide estimate, reflecting the average amount of genetic differentiation. Parameter *n* is the number of populations, *Χ*^2^_*n*-1_ is the associated chi-square value, and *n* – 1 is the associated degrees of freedom for the chi-square test (Koch, 2019). We rearranged this formula to solve *Χ*^2^_*n*-1_ for each population pair’s *Q*_ST_, for each phenotypic PC, and their respective genome-wide *F*_ST_. Associated *P*-values for *Χ*^2^_*n*-1_ values were obtained using R’s *pchisq* function. This procedure therefore tests whether estimates of additive genetic differentiation (*Q*_ST_) exceed that of the genome-wide background (*F*_ST_) (Liu & Edge, 2025).

#### Microgeographic structuring of head shape phenotypes

We tested for phenotype−environment associations between phenotypic PC axes defined at the macrogeographic scale and environmental variation among tide pools at a microgeographic scale (tens of meters) to determine whether phenotypic variation observed across broader spatial scales (hundreds of kilometres) has ecological relevance at fine spatial scales. For these analyses, we focused on fish collected at Hastings Point in 2017 and 2018, and we only analysed tide pools for which at least 3 fish had been sampled in both years (*n* = 160 and 272 fish, respectively).

We first fit models separately for each annual population using R’s *lm* function:

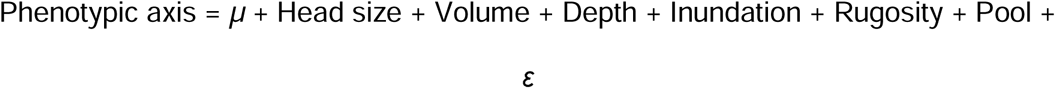

Here, the response, phenotypic axis PC1, PC2, or PC3, is fitted as a function of the continuous fixed effects of ‘Head size (log10-transformed centroid size), ‘Volume’, ‘Depth’, ‘Inundation’, and ‘Rugosity’. Each of these fixed effect predictor variables was standardised to a mean of 0 and a variance of 1. The ‘Pool’ term represents the random intercept for each tide pool, and accounts for any other, unmeasured, environmental contribution to phenotypic similarity of fish captured from the same pool. Terms *µ* and *ε* are the global mean and the residual, respectively. The significance of each fixed effect term was assessed via Type II SS ANOVA, implemented using the *Anova* function from the *car* package.

We next quantified the Spearman’s rank correlation between years for tide pools to more generally test for consistency in phenotypic structure year-to-year. We first calculated mean phenotypic PC scores per pool per year, and then partialled out the effect of mean head size in each year using R’s *lm* function:

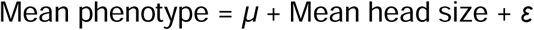

Here, the response, the tide pool mean value for phenotypic axes PC1, PC2, or PC3, is fitted as a function of the continuous effect of ‘Mean head size’, with mean *µ* and residual *ε*. The residuals from these models were retained, and the Spearman’s rank correlation coefficient between tide pools in 2017 and 2018 was estimated using the *pcor* function from R’s *ppcor* package (Kim, 2015).

## RESULTS

### Macrogeographic structuring of genotypic and phenotypic variation

Analysis of genetic variation suggested that spatiotemporally sampled *B. cocosensis* populations are subtly structured. Pairwise *F*_ST_ranged from 0.001 to 0.004 (Supplementary Table S2). These values were very low, but no randomised permutation of the data produced an *F*_ST_ greater than that observed, suggesting biological processes have structured genomic variation. Nonetheless, the PCA of genotypes did not reveal any clear groupings of populations (Supplementary Figure S2), suggesting genomic variation was predominantly within populations. Genotypic PC1 and PC2 (explaining 1.48% and 0.87% of the variation, respectively) possibly captured a chromosomal inversion, as evidenced by a 3-cluster split, a pattern associated with inversion haplotypes (Huang *et al*., 2020; Tepolt *et al*., 2021).

However, the inversion appeared to be segregating within populations and there was no clear differentiation among them (Figure 1e; Supplementary Figure S2b).

Our geometric morphometric analysis (phenotypic PC axes) allowed us to measure complex variation in head shape but was also supplemented with more tangible classic measures (distances and angles) for interpretation. For phenotypic PC1, increasing scores were associated with less angled, larger eyes and smaller mouths (Figure 2, left most column; see also Supplementary Figure S4). Phenotypic PC2 predominantly captured variation in aspects of head shape other than these three classical traits (Figure 2, centre column; see also Supplementary Figure S4).

**Figure 2.**
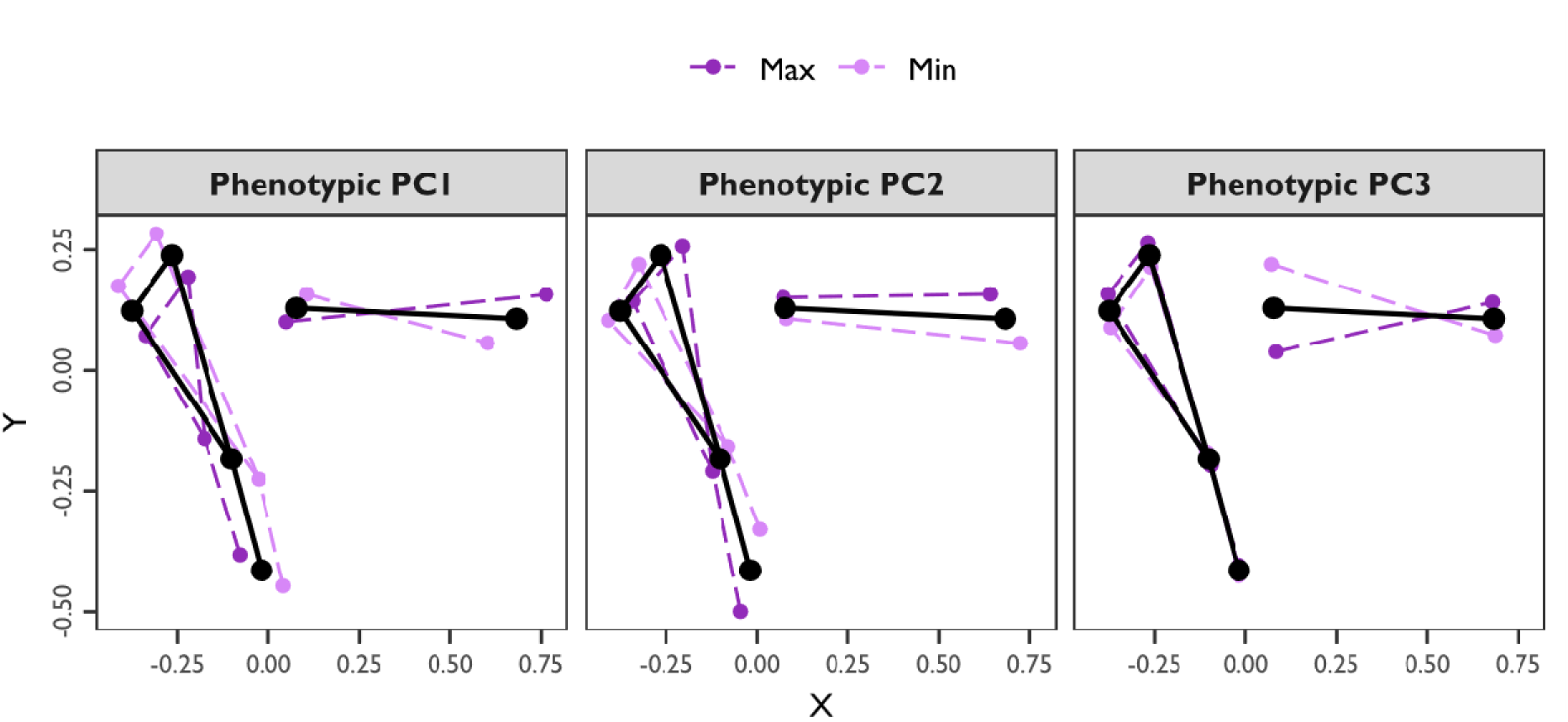
Predicted 2D head shape coordinate shifts along each phenotypic PC axis. Points indicate the measured landmark coordinates. Lines are used to provide a rough depiction of the mouth and eye shape. Large black points and solid lines are the observed mean head shape. Dark and light purple points and dashed lines are the predicted shifts in the mean shape toward the maximum (+3 SD) and minimum (−3 SD) extremes of each axis. Panels separate phenotypic axes PC1, PC2, and PC3. Predicted coordinates were estimated following: 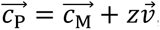, where, 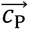, the predicted linear vector of 2D coordinates is a function of 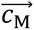, the vector of mean head shape coordinates, the desired PC axis score, *z*, and the eigenvector describing to projection of coordinates into PC space, 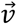. See also Supplementary Figure S2.

Phenotypic PC3 was associated with variation in eye angle and mouth size, where higher PC3 scores had less angled eyes but slightly larger mouths than fish with low PC3 scores (Figure 2, right most column; see also Supplementary Figure S4).

Pairwise *P*_ST_ values were greater than pairwise *F*_ST_ values, suggesting greater structuring of total phenotypic variation relative to genetic variation (Supplementary Figures S5). Pairwise *P*_ST_ for phenotypic PC1 ranged from 0.014 to 0.244, for PC2 it ranged from 0.015 to 0.196, and for PC3 it ranged from 0.012 to 0.470 (Supplementary Table S2). phenotypic divergence was statistically significant for many population pairs, which exhibited permuted *P* < 0.05 (Supplementary Table S2). There were no clear spatial or temporal patterns of variation in head shape: fish from the same site varied from year-to-year to a similar extent to their differentiation from other sites (Supplementary Figure S4). Decomposing the variation across all 3 head shape PCs to site and year explicitly (using MANOVA) suggested that the variance among years (11.2%) was nearly twice as much as that among sites (6.4%), with no evidence of site-specific among-year variation (site-by-year accounted for only 0.5% of the variation) (Supplementary Table S3).

### Heritability and macrogeographic structuring of additive genetic variance in head shape

Additive genetic variance, estimated across the total population of samples, was strongly supported for phenotypic PC1, with weak support for phenotypic PC2, and none for phenotypic PC3 (Table 1). Thus, our analyses suggest that *B. cocosensis* has evolutionary capacity for some aspects of head shape, particularly that associated with relative size and angle of the eye, and mouth size (Figure 2, left column).

**Table 1.** Quantitative genetic estimates from genomic animal models of head shape variation at macrogeographic scales in *B. cocosensis*. This includes variance components (*V*_F_, variance due to fixed effects; *V*_A_, additive genetic variance; and *V*_E_, residual variance), heritability (with MVN confidence intervals) and likelihood ratio tests for the significance of additive genetic effects (*Χ*^2^_*n*-1_, df and *P*-value; asterisks mark significant effects, *P* < 0.05).

| **Trait** | **$V_F$** | **$V_A$** | **$V_E$** | **$h^2$ (CI)** | **$\chi^2$** | **df** | **$P(\chi^2)$** |
| --- | --- | --- | --- | --- | --- | --- | --- |
| Pheno PC1 | 0.0012 | 0.0015 | 0.0000 | 0.57 (0.23,0.89) | 13.3 | 1 | <0.001 * |
| Pheno PC2 | 0.0014 | 0.0004 | 0.0005 | 0.17 (-0.04,0.38) | 3.2 | 1 | 0.073 |
| Pheno PC3 | 0.0003 | 0.0000 | 0.0012 | 0.00 (-0.36,0.36) | 0.0 | 1 | 1.000 |

Pairwise *Q*_ST_ values, measuring the additive genetic differentiation between populations, and were of a similar magnitude to *F*_ST_ values. Pairwise *Q*_ST_ for phenotypic PC1 ranged from 0.00002 to 0.004, for PC2 it ranged from 0.00005 to 0.002, and for PC3 it ranged from 0 to 0.001 (Supplementary Table S2). Chi-square tests (df = 1 = *n* populations – 1) failed to reject the null hypothesis that *Q*_ST_ = *F*_ST_ (*α* = 0.05). These results suggest that additive genetic differentiation among population pairs is not greater than expected under neutral evolution (genetic drift). Thus, despite evidence that there is additive genetic variance in some dimensions of head shape, there is no statistical evidence that the heritable component is due to adaptive divergence in phenotypes.

### Microgeographic structuring of head shape phenotypes

Within Hastings Point, there was some evidence that tide pool environment explained head shape variation, but phenotype-environment associations were not temporally stable (Table 2). In the 2017 population, both phenotypic PC2 and PC3 scores increased as inundation time decreased (Figure 3; Supplementary Table S4). However, in the 2018 population, inundation time influenced only PC3, and in the opposite direction to 2017 (Figure 3; Supplementary Table S4). In 2018, generally, there were stronger associations of environmental factors and phenotypic PCs: phenotypic PC1 scores were positively associated with depth and a negatively associated with volume; PC2 exhibited a positive association with depth, volume and rugosity; PC3 exhibited a negative association with depth and rugosity in addition to the positive association with inundation (Figure 3; Supplementary Table S4). As expected from these temporally unstable environmental associations, Spearman rank correlations in mean phenotypic PC scores (after accounting for size) were low and not statistically distinguishable from zero (PC1: *r* = −0.15, *P* = 0.67; PC2: *r* = 0.38, *P* = 0.27; and PC3: *r* = 0.19, *P* = 0.60).

**Figure 3.**
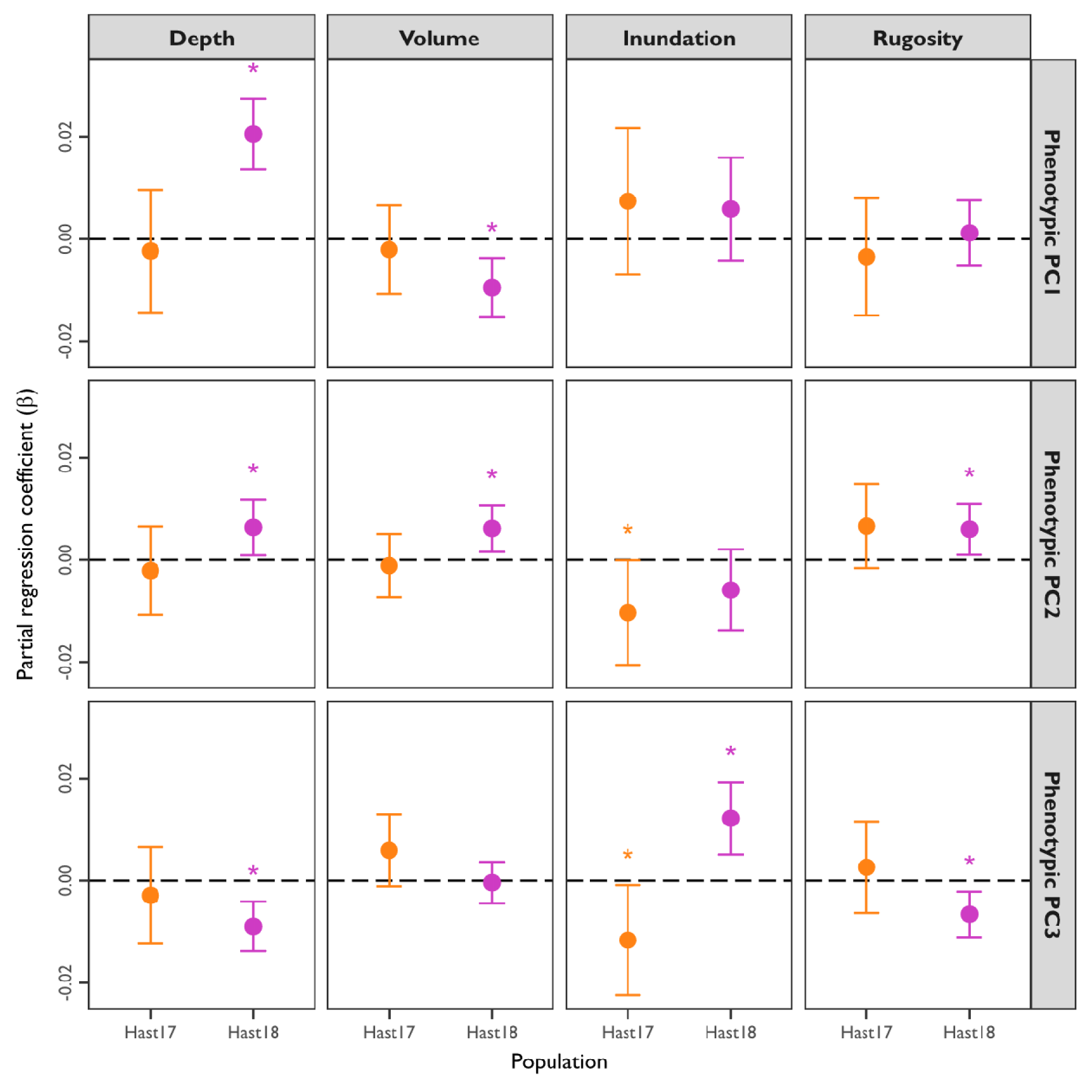
Phenotype−environment associations at microgeographic scales in *B. cocosensis* from Hastings Point. The *x*-axis represents populations, and the *y*-axis represents the partial regression coefficient estimates. Points represent estimates for each population with 95% confidence intervals. Asterisks above points indicate significance (*P* < 0.05) in an analysis of deviance (see Table 2). Dashed lines indicate coefficient estimates of 0. Panels separate data for different combinations of phenotypic PC axis (in rows) and predictor variables (head size and microhabitat; in columns).

**Table 2.** Analysis of deviance for phenotype−environment associations at microgeographic scales for *B. cocosensis* at Hastings Point. Asterisks mark significant effects (*P* < 0.05).

| **Population** | **Trait** | **Predictor** | **$\chi^2$** | **df** | **$P(\chi^2)$** |
| --- | --- | --- | --- | --- | --- |
| Hast17 | PC1 | Head size | 83.52 | 1 | <0.001 * |
|  |  | Depth | 0.15 | 1 | 0.700 |
|  |  | Volume | 0.22 | 1 | 0.638 |
|  |  | Inundation | 1.04 | 1 | 0.309 |
|  |  | Rugosity | 0.36 | 1 | 0.551 |
|  | PC2 | Head size | 164.19 | 1 | <0.001 * |
|  |  | Depth | 0.23 | 1 | 0.631 |
|  |  | Volume | 0.13 | 1 | 0.718 |
|  |  | Inundation | 3.90 | 1 | 0.048 * |
|  |  | Rugosity | 2.47 | 1 | 0.116 |
|  | PC3 | Head size | 28.70 | 1 | <0.001 * |
|  |  | Depth | 0.35 | 1 | 0.551 |
|  |  | Volume | 2.73 | 1 | 0.098 |
|  |  | Inundation | 4.49 | 1 | 0.034 * |
|  |  | Rugosity | 0.33 | 1 | 0.568 |
| Hast18 | PC1 | Head size | 31.75 | 1 | <0.001 * |
|  |  | Depth | 33.84 | 1 | <0.001 * |
|  |  | Volume | 10.41 | 1 | 0.001 * |
|  |  | Inundation | 1.30 | 1 | 0.254 |
|  |  | Rugosity | 0.14 | 1 | 0.708 |
|  | PC2 | Head size | 138.89 | 1 | <0.001 * |
|  |  | Depth | 5.16 | 1 | 0.023 * |
|  |  | Volume | 6.90 | 1 | 0.009 * |
|  |  | Inundation | 2.12 | 1 | 0.146 |
|  |  | Rugosity | 5.46 | 1 | 0.019 * |
|  | PC3 | Head size | 18.55 | 1 | <0.001 * |
|  |  | Depth | 13.05 | 1 | <0.001 * |
|  |  | Volume | 0.03 | 1 | 0.853 |
|  |  | Inundation | 11.46 | 1 | 0.001 * |
|  |  | Rugosity | 8.40 | 1 | 0.004 * |

## DISCUSSION

In this study, we provide one of the most comprehensive comparisons of genetic and phenotypic variation in a wild marine organism that exhibits high effective gene flow, large population sizes, and with limited feasibility to sample close relatives: *Bathygobius cocosensis*. We also we provide the first estimates of heritability in a marine cryptobenthic fish, demonstrating the value of genomic animal models to derive quantitative genetic estimates in wild marine organisms. We found that spatial structuring of head shape phenotypes, across broad macrogeographic scales and multiple years, exceeds the level of genomic and quantitative genetic differentiation. These results suggest that much of the variation in head shape at broad spatiotemporal scales is likely due to non-heritable plastic variation. Our microgeographic scale interrogation of phenotype–environment associations did not identify specific factors leading to consistent structuring of head shape variation. The relative importance of heritable versus plastic responses to local microhabitat heterogeneity remains a scope for future work.

### Additive genetic variance in head shape

A key finding of our present work was that head shape phenotypes in *B. cocosensis* have a heritable component. Estimated narrow-sense heritability for phenotypic PC1, ℎ^2^ *=* 0.54 (LRT *P* < 0.001), is consistent with published estimates for morphological traits in wild populations of terrestrial taxa: mean ℎ^2^ = 0.56 (±0.035 SE) (reviewed in Postma 2014). There was weak statistical support for additive genetic variance in phenotypic PC2, with estimated ℎ^2^ *=* 0.17 (LRT *P* = 0.073), and no evidence for phenotypic PC3, with estimated ℎ^2^ *=* 0.00 (LRT *P* = 1). It is possible that our estimates may be deflated relative to some published studies because we included the phenotypic variance attributed to fixed effects in our estimate of ℎ^2^ (Wilson *et al*., 2010b; de Villemereuil *et al*., 2018). For marine taxa with large populations sizes, like *B. cocosensis*, obtaining a kin structured dataset can be very challenging, even if many individuals can be sampled from the wild (Frentiu *et al*., 2008; Fraimout *et al*., 2024). However, the strong signal of heritability in some components of *B. cocosensis* head shape highlights the potential of genomic animal models for advancing our understanding of ecologically important morphological traits in marine taxa from realistic sample sizes.

### Phenotypic differentiation at macrogeographic scales exceeds that of genetic differentiation

Our present study indicates that there is broad-scale genetic homogeneity in *B. cocosensis*, which is in line with previous work on different populations (Thia *et al*., 2021). Our estimates of genetic differentiation across macrogeographic scales were very subtle across distant sites (∼600 km) and through time (2 years), with 0.001 ≤ *F*_ST_ ≤ 0.004. Like many marine organisms, *B. cocosensis* has a planktonically dispersing larval stage that may afford long-distance dispersal (Thia *et al*., 2018), which is typically expected to homogenise genetic variation over broad spatial scales and over generations (Waples, 1998; Babin *et al*., 2017; Haro-Bilbao *et al*., 2021).

However, processes like sweepstakes reproduction (Hedgecock & Pudovkin, 2011), variable dispersal (D’Aloia & Neubert, 2018), and heterogeneous selection (Schmidt & Rand, 1999), can cause low levels of transient and patchy structuring of genetic variation in marine systems – so called ‘chaotic genetic patchiness’ (Johnson & Black, 1982). Such processes have previously been hypothesised and tested in *B. cocosensis* across its range on Australia’s east coast, potentially contributing to general patterns of weak spatiotemporal genetic structure in this cryptobenthic marine fish (Thia *et al*., 2021).

Our genomic animal models allowed us to perform classic *F*_ST_ versus *Q*_ST_ comparisons as a test of adaptive genetic divergence among populations. These tests are inaccessible when estimates of the additive genetic variance among and within populations are unavailable (Merilä & Crnokrak, 2001). Genomic animal models thus provide great utility in this regard by facilitating genomically informed estimates of additive genetic breeding values, which can then be used to calculate *Q*_ST_. In this study of *B. cocosensis*, we found that pairwise estimates of additive genetic differentiation among populations (based on additive genetic breeding values) did not exceed expectations from the genome-wide background (Liu & Edge, 2025), with 0.00001 ≤ *Q*_ST_ ≤ 0.004. Accordingly, additive genetic variance in head shape appears to be distributed primarily within, rather than among, populations across macrogeographic scales and over time. That is, our results do not suggest local adaptation in head shape due to genetic divergence (Merilä & Crnokrak, 2001).

In contrast to this low genomic and quantitative genetic structure, phenotypic variation in head shape, quantified along three principal component axes derived from morphometric landmarks, was generally greater, 0.0001 ≤ *P*_ST_ ≤ 0.47. In systems with high gene flow, where individuals are likely to encounter different environments to their parents, phenotypic plasticity is predicted to evolve as an adaptive mechanism to match phenotype to environment (Scheiner, 2013). In fishes, head morphology is closely linked to ecologically important functions, including prey acquisition and sensory performance, such as vision for navigation and predator avoidance (Winemiller *et al*., 1995; Willis *et al*., 2005; Caves *et al*., 2017; Vera-Duarte *et al*., 2017; Andersson *et al*., 2024). Consistent with this, our current findings, and those from previous work (Thia *et al*., 2021), suggest that environmental differences may drive phenotypic differentiation across macrogeographic scales and among years in *B. cocosensis*. Notably, the phenotypic differentiation over time (between years) was greater than between the sites, suggesting that temporal heterogeneity in environmental conditions may play an important role in generating phenotypic variation in this system.

### Phenotype−environment associations on microgeographic scales are temporally variable

Our results suggest that multiple aspects of head shape (such as eye angle, eye shape and mouth size) are associated with physical microhabitat variables (volume, depth, inundation time, and rugosity). Some of the most detailed work on microhabitat structuring of phenotypic variation in intertidal zones has come from studies on barnacles and gastropods (Schmidt & Rand, 1999; Johnson & Black, 2008; Phifer-Rixey *et al*., 2008; Westram *et al*., 2014). However, these organisms have much lower adult mobility than fish and may be constrained in the environments they are exposed to once they have settled on the benthos. In contrast, the greater mobility of intertidal fish can allow them to move away from unsuitable microhabitats or preferentially occupy suitable ones. Nonetheless, *B. cocosensis* exhibits strong homing behaviour, returning to specific tide pools at low tide or after displacement (Griffiths, 2003a; White & Brown, 2013; Malard et al., 2016). This site fidelity may expose individuals to consistent microhabitat conditions over time, reinforcing phenotype–environment associations despite their mobility.

Although we found significant associations of each phenotypic PC axis with environmental factors, these associations were not consistent across populations sampled in 2017 and 2018 at Hastings Point (Figure 3). Local temporal variability in environmental selection has been reported in other studies (Stratton & Bennington, 1998; Mojica *et al*., 2012; Rudman *et al*., 2022). It remains to be determined whether the observed differences reflect variable effects of tide pool environmental factors (either those measured and analysed here, or other, unmeasured, characteristics), or instead reflect earlier, pre-settlement heterogeneity in environmental condition.

Further, while we could consider the genomic and quantitative genetic structuring among tide pools, the relatively small sample sizes per pool preclude accurate estimation. This leaves open the question of whether, on this microgeographic scale, genetic differentiation plays a different role than that observed at the macrogeographic scale.

## CONCLUDING REMARKS

In summary, our study shows that phenotypic differentiation in head shape in *Bathygobius cocosensis* can exceed genetic differentiation across large spatial scales and among annual cohorts. Although we detected heritable variation in some aspects of head shape morphology, this variation does not show clear signatures of adaptive genetic divergence at macrogeographic scales or between temporally discrete populations. This pattern is consistent with a prominent role for phenotypic plasticity in shaping observed differentiation, highlighting the potential for flexible responses to environmental variation. Future work examining how head shape develops under contrasting environmental conditions, and how this variation relates to fitness, will be key to understanding its ecological and evolutionary significance. Coupled with more detailed investigation of complex environmental parameters, this may allow us to develop clearer insight into the causal environmental factors.

Overall, our work highlights the value of integrating quantitative genomic approaches with ecological measurements to investigate the processes shaping phenotypic variation in non-model organisms, including marine taxa. Broader uptake of quantitative genomics in evolutionary studies would enable more rigorous testing of hypotheses about the role of adaptation in driving phenotypic divergence.

## ACKNOWLEDGEMENTS

We thank I. Popovic, C. Da Silva, A. Matias, A. Mather, and D. Blower for field assistance. We thank V. Paris, O. Holland, J. Chang and J. Brown for providing feedback on this manuscript. Fish were collected under NSW DPI Permit P13/0046-1.1, and from QLD under the QLD Government Permit #174684. Funding was provided by The Hermon Slade Foundation (HSF 13/14 to CR and LL) and The Ecological Society of Australia (Student Research Award 2015 and the Wiley Fundamental Ecology Award 2016 to JAT).

## AUTHOR CONTRIBUTIONS

Design of study: co-led by JAT, KM, CR; contributions by JE. Sample collection: co-led by JAT and JE. Genetic data collection: led by JAT; contributions by JH. Phenotype data collection: co-led by JAT and JE. Statistical analysis: led by JAT; contributions by KM and JDA. Archiving data: led by JAT. Funding: contributions by CR, LL, and JAT. Permits: led by CR and WF. Supervision: CR, KM, and LL. Drafting of original manuscript: JAT. Comments on final draft: all authors.

## DATA AVAILABILITY

All data and scripts associated with this study have been uploaded to a University of Melbourne repository hosted by FigShare: doi.org/10.26188/32684940. This dataset will be made publicly available upon publication of this manuscript.

## AI USE STATEMENT

ChatGPT v5.2 was used by JAT to curate some of the code for this study. This included assistance in deriving calculations, increasing the efficiency of complex pipelines, and building of custom functions.

## SUPPLEMENTARY FIGURES

**Figure S1.**
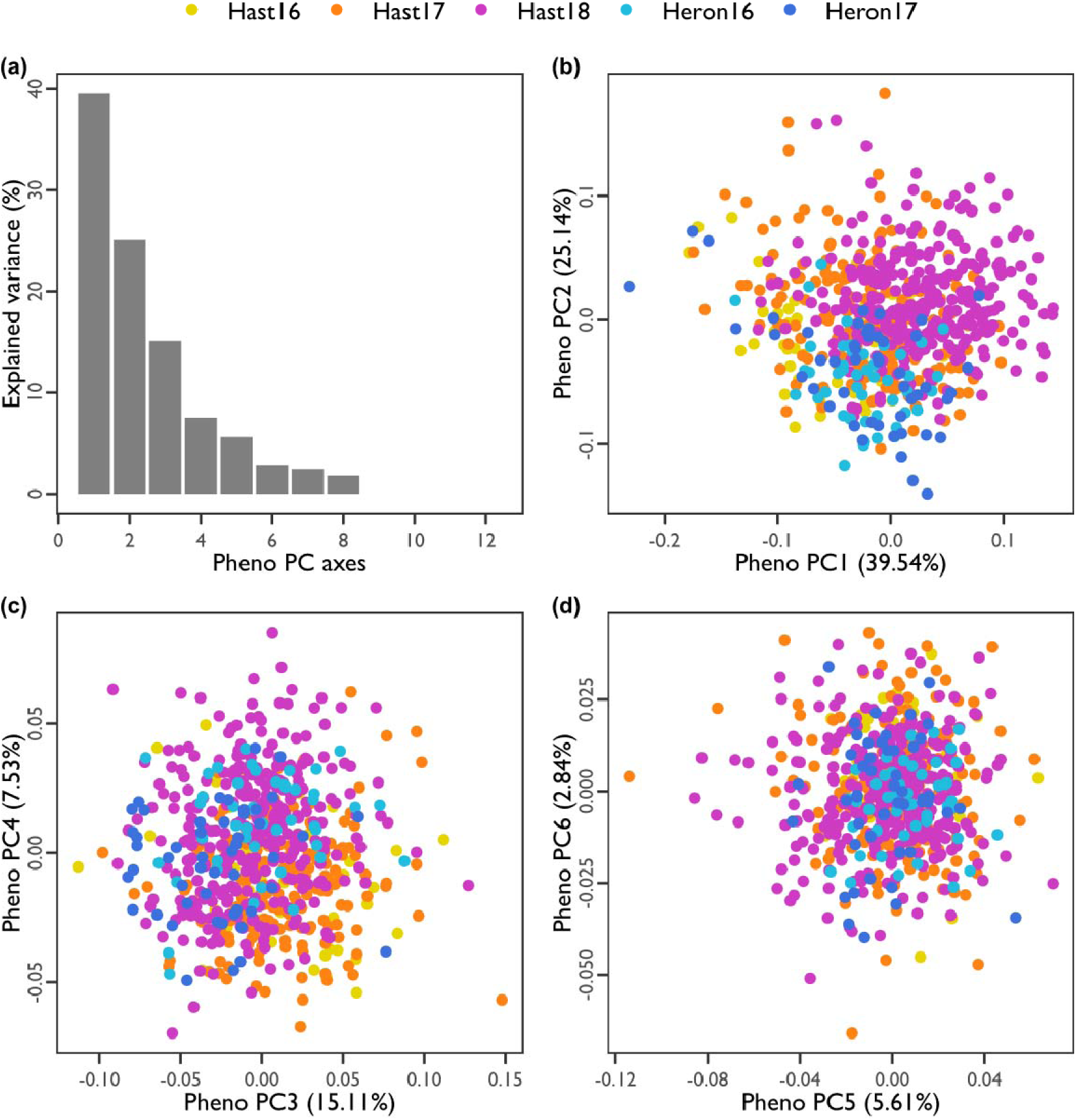
PCA of phenotypic variation in *B. cocosensis*. (a) Screeplot of explained variance for the first 20 PC axes. The *x*-axis represents the PC axes, and the *y*-axis is the precent of the total variance explained by each axis. The leading 3 PCs explained more phenotypic variance than expected by sampling error at *α* = 0.05. (b) Scatterplot of PC1 vs PC2. (c) Scatterplot of PC3 vs PC4). (d) Scatterplot of PC5 vs PC6. (a–d) Points represent PC scores of individual fish, coloured by population (see legend).

**Figure S2.**
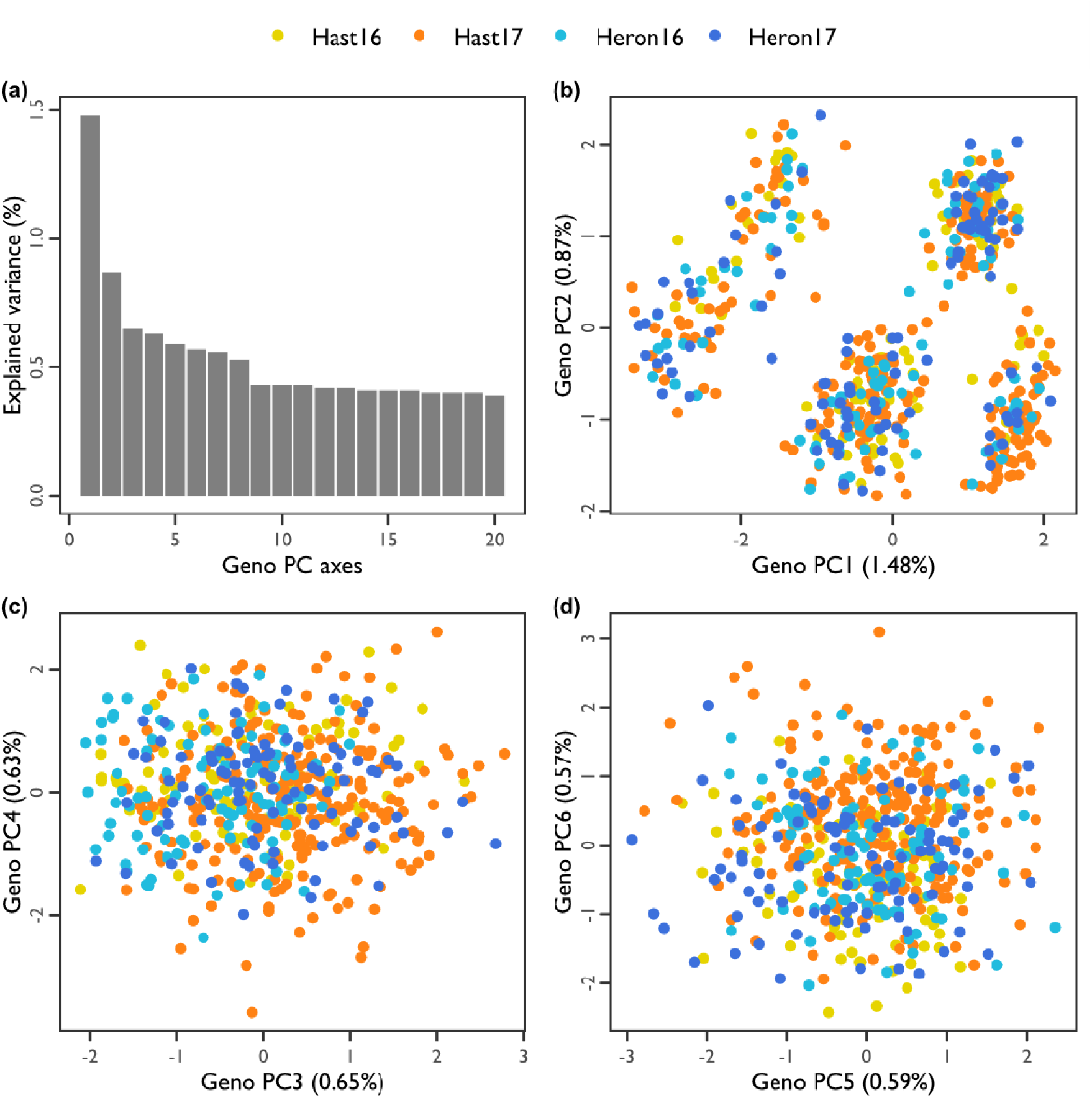
PCA of genotypic variation in *B. cocosensis*. (a) Screeplot of explained variance for the first 20 PC axes. The *x*-axis represents the PC axes, and the *y*-axis is the precent of the total variance explained by each axis. Randomised permutation indicated that the leading 18 genotypic PC axes accounted for significantly more variation than expected from sampling error alone (at *α* = 0.05). (b) Scatterplot of PC1 vs PC2. (c) Scatterplot of PC3 vs PC4). (d) Scatterplot of PC5 vs PC6. (b–d) Points represent genomic PC scores for individual fish, coloured by population (see legend).

**Figure S3.**
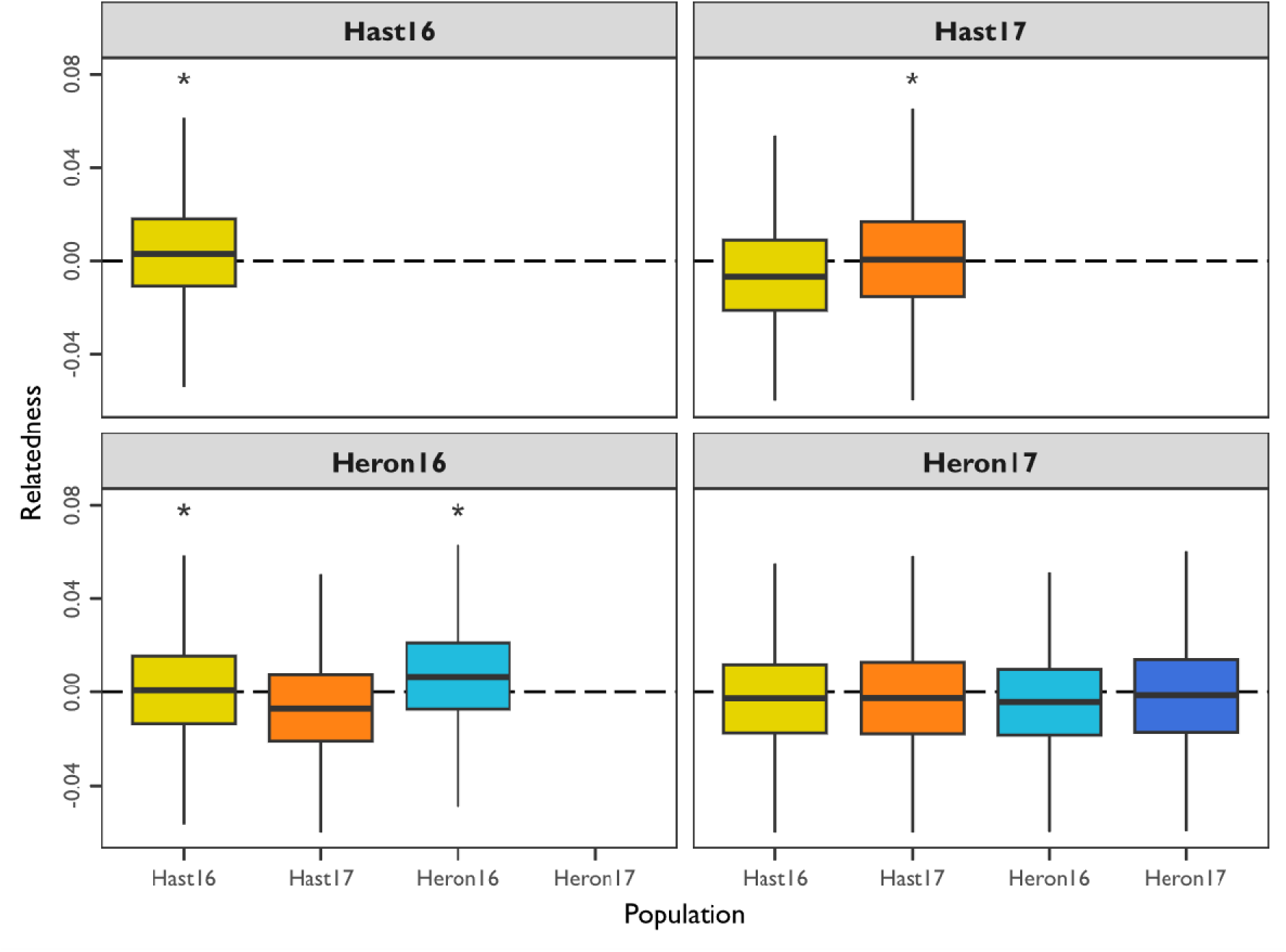
Relatedness within and between populations. Each panel shows results for a focal population. The *x*-axis denotes the contrast population used for comparison, and the *y*-axis shows the relatedness between pairs of individuals drawn from the focal and contrast populations. When the focal and contrast populations are identical, the values represent within-population relatedness; when they differ, the values represent between-population relatedness. Box plots summarise the distribution of relatedness values. Asterisks denote comparisons where the mean relatedness is greater than zero, as estimated from a single-sample *t*-test with *α* = 0.05. Expected relatedness values for different close-kin relationships: full-siblings or parent−offspring, 0.5; half-siblings, 0.25, cousins, 0.125; half-cousins, 0.0625. Our data are thus consistent with most individuals being largely unrelated.

**Figure S4.**
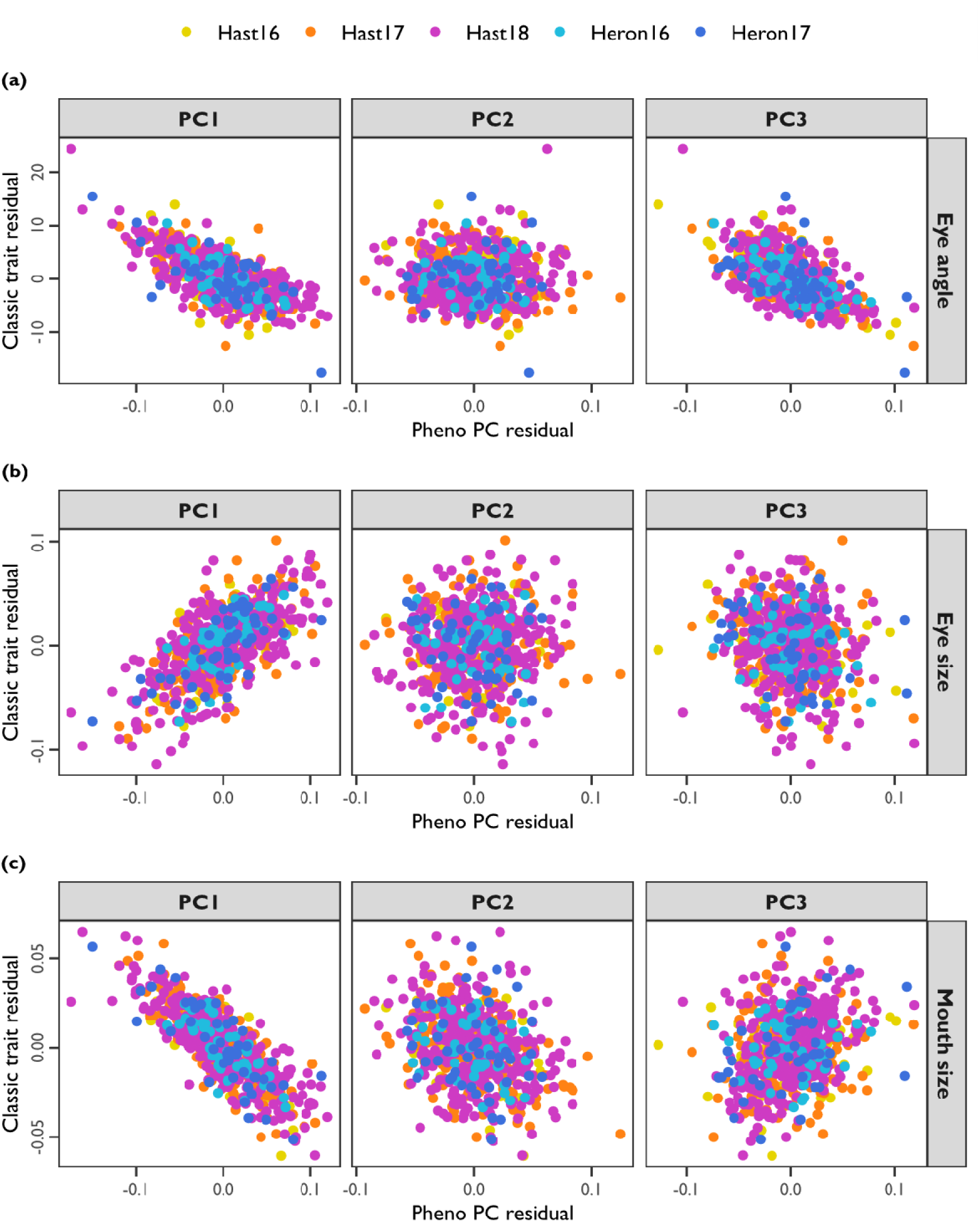
Associations between geometric and classic morphometric traits in the analysis of macrogeographic variation across Hastings Point and Heron Island. Linear models were fit to each trait (PC or classic) to account for variation due to head size and population, and the residuals are plotted here (PCs on *x*-axes and classic trait on *y*-axes). Points represent individual fish, coloured by population (see legend). Panels contain data for each phenotypic PC (columns). Classically measured traits included: (a) eye angle, (b) eye size, and (c) mouth size.

**Figure S5.**
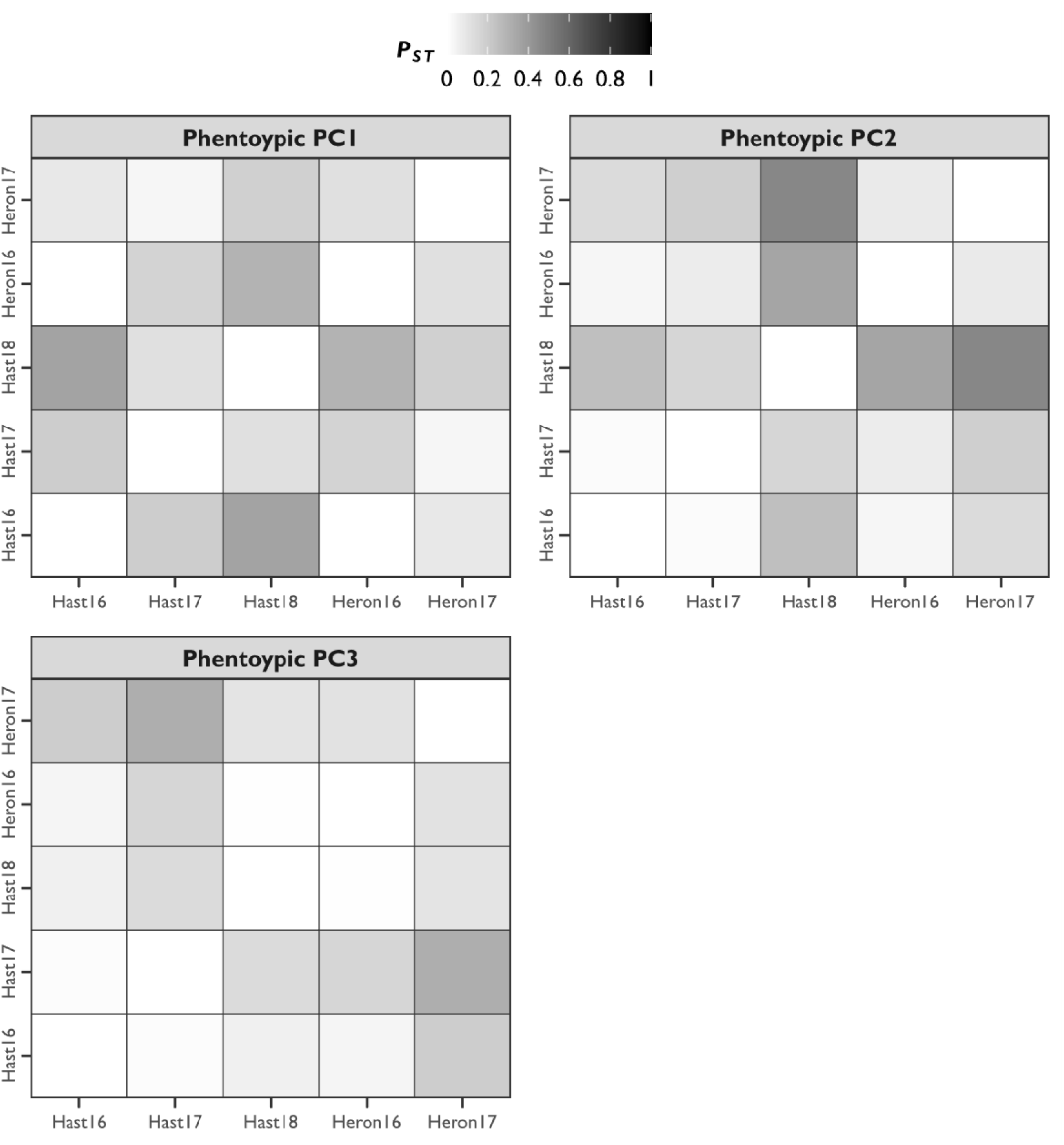
Heatmaps of pairwise *P*_ST_ values. Populations are arranged on the *x*- and *y*-axes. Shading of cells indicates the magnitude of *P*_ST_ (see legend). Panels separate estimates for phenotypic axes PC1, PC2, and PC3.

## SUPPLEMENTARY TABLES

**Table S1.** Analysis of variance for the relationship between phentoypic PC axes and classic traits (after partialling out the effect of head size and population)

| **Phenotypic axis** | **Classic trait** | **Term** | **Sum Sq** | **Df** | **F** | **P(F)** |
| --- | --- | --- | --- | --- | --- | --- |
| PC1 | Eye angle | Axis | 5820.965 | 1 | 571.492 | 2.54E-91 |
|  |  | Residuals | 6681.726 | 656 |  |  |
|  | Eye size | Axis | 0.320064 | 1 | 494.1865 | 4.88E-82 |
|  |  | Residuals | 0.424864 | 656 |  |  |
|  | Mouth size | Axis | 0.18736 | 1 | 1373.741 | 4.81E-163 |
|  |  | Residuals | 0.089469 | 656 |  |  |
| PC2 | Eye angle | Axis | 2.04354 | 1 | 0.107239 | 0.743413657 |
|  |  | Residuals | 12500.65 | 656 |  |  |
|  | Eye size | Axis | 1.90E-05 | 1 | 0.016718 | 0.897161985 |
|  |  | Residuals | 0.744909 | 656 |  |  |
|  | Mouth size | Axis | 0.031811 | 1 | 85.17077 | 3.71E-19 |
|  |  | Residuals | 0.245018 | 656 |  |  |
| PC3 | Eye angle | Axis | 4616.156 | 1 | 383.9707 | 1.18E-67 |
|  |  | Residuals | 7886.535 | 656 |  |  |
|  | Eye size | Axis | 0.023031 | 1 | 20.92881 | 5.70E-06 |
|  |  | Residuals | 0.721897 | 656 |  |  |
|  | Mouth size | Axis | 0.01671 | 1 | 42.14031 | 1.68E-10 |
|  |  | Residuals | 0.260119 | 656 |  |  |

**Table S2.** Pairwise genetic, phenotypic, and additive genetic differentiation at macrogeographic scales in *B. cocosensis*.

| **Trait** | **Pop1** | **Pop2** | **Fst** | **P(Fst)** | **Pst** | **P(Pst)** | **Qst** | **P(Qst)** |
| --- | --- | --- | --- | --- | --- | --- | --- | --- |
| PC1 | Hast16 | Hast17 | 0.003 | 0.000 | 0.197 | 0.000 | 0.004 | 0.701 |
|  | Hast16 | Heron16 | 0.002 | 0.000 | 0.000 | 0.892 | 0.000 | 0.073 |
|  | Hast16 | Heron17 | 0.002 | 0.000 | 0.090 | 0.008 | 0.000 | 0.144 |
|  | Hast17 | Heron16 | 0.004 | 0.000 | 0.170 | 0.000 | 0.003 | 0.626 |
|  | Hast17 | Heron17 | 0.001 | 0.000 | 0.033 | 0.024 | 0.003 | 0.895 |
|  | Hast18 | Hast16 |  |  | 0.361 | 0.000 |  |  |
|  | Hast18 | Hast17 |  |  | 0.121 | 0.000 |  |  |
|  | Hast18 | Heron16 |  |  | 0.296 | 0.000 |  |  |
|  | Hast18 | Heron17 |  |  | 0.185 | 0.000 |  |  |
|  | Heron16 | Heron17 | 0.004 | 0.000 | 0.119 | 0.000 | 0.000 | 0.066 |
| PC2 | Hast16 | Hast17 | 0.003 | 0.000 | 0.016 | 0.112 | 0.002 | 0.547 |
|  | Hast16 | Heron16 | 0.002 | 0.000 | 0.033 | 0.052 | 0.001 | 0.478 |
|  | Hast16 | Heron17 | 0.002 | 0.000 | 0.139 | 0.000 | 0.000 | 0.128 |
|  | Hast17 | Heron16 | 0.004 | 0.000 | 0.074 | 0.000 | 0.000 | 0.204 |
|  | Hast17 | Heron17 | 0.001 | 0.000 | 0.187 | 0.000 | 0.002 | 0.892 |
|  | Hast18 | Hast16 |  |  | 0.251 | 0.000 |  |  |
|  | Hast18 | Hast17 |  |  | 0.161 | 0.000 |  |  |
|  | Hast18 | Heron16 |  |  | 0.353 | 0.000 |  |  |
|  | Hast18 | Heron17 |  |  | 0.469 | 0.000 |  |  |
|  | Heron16 | Heron17 | 0.004 | 0.000 | 0.080 | 0.008 | 0.001 | 0.432 |
| PC3 | Hast16 | Hast17 | 0.003 | 0.000 | 0.010 | 0.132 | 0.000 |  |
|  | Hast16 | Heron16 | 0.002 | 0.000 | 0.039 | 0.052 | 0.000 |  |
|  | Hast16 | Heron17 | 0.002 | 0.000 | 0.195 | 0.000 | 0.000 |  |
|  | Hast17 | Heron16 | 0.004 | 0.000 | 0.158 | 0.000 | 0.000 |  |
|  | Hast17 | Heron17 | 0.001 | 0.000 | 0.318 | 0.000 | 0.000 |  |
|  | Hast18 | Hast16 |  |  | 0.058 | 0.000 |  |  |
|  | Hast18 | Hast17 |  |  | 0.141 | 0.000 |  |  |
|  | Hast18 | Heron16 |  |  | 0.003 | 0.420 |  |  |
|  | Hast18 | Heron17 |  |  | 0.103 | 0.000 |  |  |
|  | Heron16 | Heron17 | 0.004 | 0.000 | 0.114 | 0.000 | 0.000 |  |

**Table S3.**
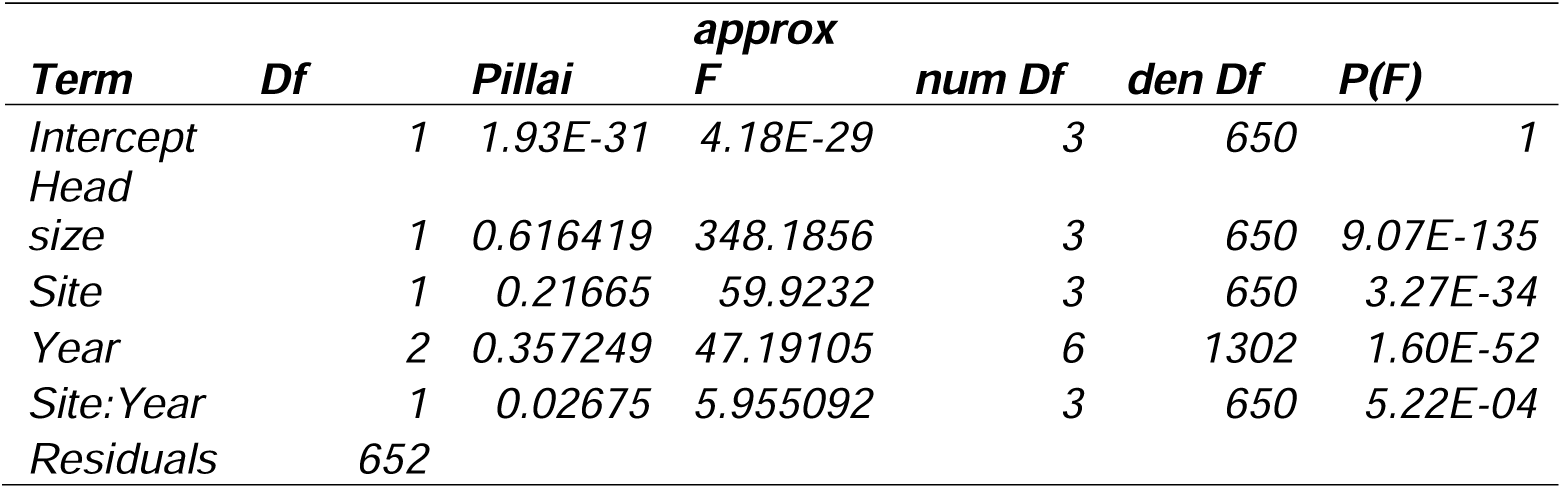
Multivariate analysis of variance of phenotypic PC1, PC2 and PC3.

| **Term** | **Df** | **Pillai** | **approx**<br>**F** | **num Df** | **den Df** | **P(F)** |
| --- | --- | --- | --- | --- | --- | --- |
| Intercept | 1 | 1.93E-31 | 4.18E-29 | 3 | 650 | 1 |
| Head<br>size | 1 | 0.616419 | 348.1856 | 3 | 650 | 9.07E-135 |
| Site | 1 | 0.21665 | 59.9232 | 3 | 650 | 3.27E-34 |
| Year | 2 | 0.357249 | 47.19105 | 6 | 1302 | 1.60E-52 |
| Site:Year | 1 | 0.02675 | 5.955092 | 3 | 650 | 5.22E-04 |
| Residuals | 652 |  |  |  |  |  |

**Table S4.** Partial regression coefficients for models of phenotype−environment associations at macrogeographic scales for *B. cocosensis* at Hastings Point.

| **Year** | **Coeff** | **Estimate** | **SE** | **Z-value** | **P(Z)** |
| --- | --- | --- | --- | --- | --- |
| 2017 | *Intercept* | -0.12 | 0.1 | -1.14 | 0.252 |
|  | *Volume* | 0.33 | 0.1 | 3.18 | 0.001 |
|  | *Depth* | -0.34 | 0.14 | -2.42 | 0.016 |
|  | *Inundation* | -0.57 | 0.17 | -3.46 | 0.001 |
|  | *Rugosity* | 0.09 | 0.14 | 0.65 | 0.514 |
| 2018 | *Intercept* | -0.03 | 0.11 | -0.29 | 0.774 |
|  | *Volume* | 0.24 | 0.13 | 1.81 | 0.071 |
|  | *Depth* | -0.82 | 0.15 | -5.43 | <0.001 |
|  | *Inundation* | -0.22 | 0.18 | -1.26 | 0.209 |
|  | *Rugosity* | -0.15 | 0.15 | -0.98 | 0.329 |

